# A tRNA-gRNA multiplexing system for CRISPR genome editing in *Marchantia polymorpha*

**DOI:** 10.1101/2025.04.18.649274

**Authors:** Eftychios Frangedakis, Nataliya E. Yelina, Satish Kumar Eeda, Facundo Romani, Alexandros Fragkidis, Jim Haseloff, Julian M. Hibberd

## Abstract

The liverwort *Marchantia polymorpha* is a widely used model organism for studying land plant biology, which has also proven to be a promising testbed for bioengineering. CRISPR/Cas9 technology has emerged as a transformative tool for precise genome modifications in *M. polymorpha*. However, a robust method for the simultaneous expression of multiple gRNAs, which is crucial for enhancing the efficiency and versatility of CRISPR/Cas9-based genome editing, has yet to be fully developed. In this study, we introduce an adaptation from the OpenPlant kit CRISPR/Cas9 tools, that facilitates expression of multiple gRNAs from a single transcript through incorporation of tRNA sequences. This approach significantly improves the efficiency and scalability of genome editing in *M. polymorpha*. Additionally, by combining this vector system with a simplified and optimized protocol for thallus transformation, we further streamline the generation of CRISPR/Cas9 mutants in *M. polymorpha*. The resulting gene-editing system offers a versatile, time-saving and straightforward tool for advancing functional genomics in *M. polymorpha*, enabling more comprehensive genetic modifications and genome engineering.

## Introduction

*Marchantia polymorpha* is a powerful model organism for investigating various aspects of plant biology, including genetics, cell biology, development, and evolution (Naramoto et al. 2022; Bowman et al. 2022; Kohchi et al. 2021). *M. polymorpha* is also an excellent chassis for bioengineering, offering a unique combination of well-established genetic modification tools, simple body plan, rapid growth and remarkable regenerative capacity, as well as minimal cultivation requirements (Boehm et al. 2017; Sauret-Güeto et al. 2020; Tse et al. 2024; Frangedakis et al. 2021; Romani et al. 2024; Ishizaki et al. 2016). All these make it an ideal platform for a wide range of synthetic biology applications, such as reprogramming cell fate, engineering of biosynthetic pathways, development of biosensors, Clustered Regularly Interspaced Short Palindromic Repeats (CRISPR)-based DNA-recording device or CRISPR based multiplex gene regulation systems (Donà et al. 2023; Tansley, Patron, and Guiziou 2024; Forestier et al. 2025; Gupta and Karkute 2021; Kocaoglan, Radhakrishnan, and Nakayama 2023).

*M. polymorpha’s* compact genome and haploid dominant life cycle make it particularly well-suited for studying genetics (Kohchi et al. 2021). As a result, it is uniquely positioned to explore fundamental genetic processes, with potential to enhance our understanding of key traits that are conserved across land plants. Genetics has been profoundly transformed by the development of CRISPR technology. CRISPR enables precise and targeted genetic modifications by inducing deletions or insertions at specific loci within the nuclear genome (Ran et al. 2013; Jinek et al. 2012; Cong et al. 2013). This gene-editing system utilizes a guide RNA (gRNA) to direct a nuclease protein such as Cas9, to a specific DNA sequence, where it induces a double-strand break, allowing for accurate genome editing (Adli 2018; Ran et al. 2013). CRISPR has facilitated significant advances in functional genomics, where the precise manipulation of sequence allows a better understanding of gene function. CRISPR/Cas9 technology is routinely used in *M. polymorpha* (Sugano et al. 2018, 2014; Sauret-Güeto et al. 2020). One of the challenges in CRISPR-based technologies, including in *M. polymorpha*, is the expression of multiple guide RNAs (gRNAs) as a single transcript, particularly for applications requiring simultaneous editing of multiple genomic sites. Current CRISPR/Cas9 systems in *M. polymorpha* enable only limited multiplex genome editing or require multiple plasmids with different antibiotic resistance, typically allowing the expression of no more than two gRNAs simultaneously (Sugano et al. 2018, 2014; Sauret-Güeto et al. 2020). Expressing multiple gRNAs as part of a tRNA-gRNA transcript is a gRNA multiplexing approach with proven value in several plant species (Xie, Minkenberg, and Yang 2015; Hui et al. 2019; Z. Wang et al. 2021). In this system (**Figure A**), tRNA-gRNA synthetic transcripts mimic native tRNAsnoRNA43 transcripts in plants, allowing endogenous cellular machinery (i.e. RNase P and Z) to cleave the tRNA structure and release mature functional single gRNAs (**Figure 1B**) (Xie, Minkenberg, and Yang 2015; Ma et al. 2019; Čermák et al. 2017). Importantly, tRNA-gRNA system reduces the complexity associated with constructing multiple independent gRNA expression cassettes.

**Figure 1:**
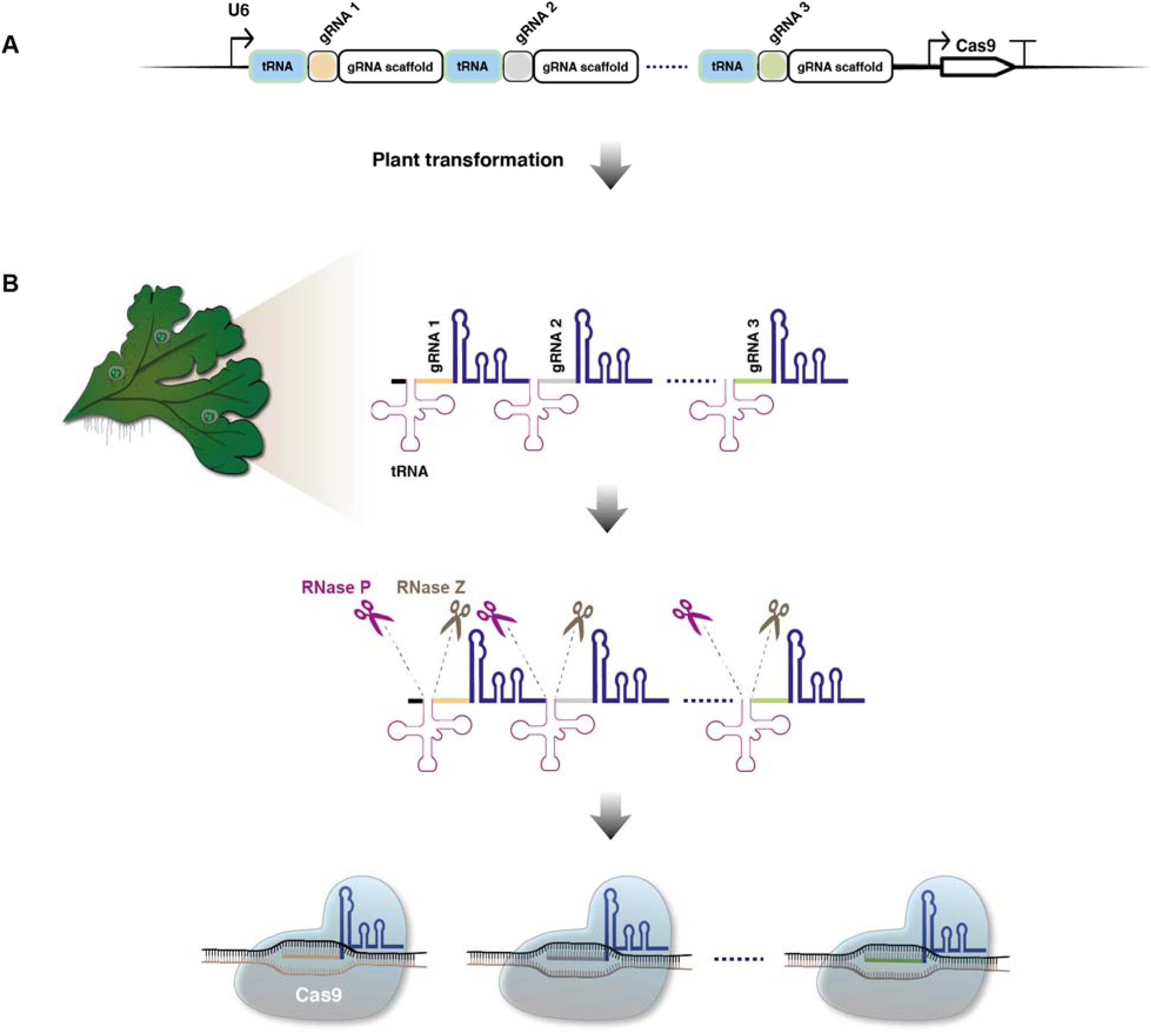
Schematic diagram of tRNA-gRNA multiplexing for plant genome editing. **(A)** A plasmid is constructed containing a tRNA-gRNA cassette driven by the U6 promoter, along with a transcription unit for Cas9 expression. **(B)** After transformation into the plant, the primary RNA transcript is cleaved by endogenous tRNA-processing enzymes, RNase P and RNase Z, resulting in functional guide RNAs for genome editing

Previously, we have generated vectors as part of the OpenPlant kit (Sauret-Güeto et al. 2020) that allow the introduction of both the Cas9 protein and up to two single gRNA expression cassette into the *M. polymorpha* genome via *Agrobacterium*-mediated plant transformation. In this study, we present an updated CRISPR/Cas9 vector system for *M. polymorpha*, adapted from the OpenPlant kit, that combines the Cas9 protein with a tRNA-gRNA module as a single transcript. Furthermore, we combine the updated CRISPR/Cas9 vector system with an optimized and simplified protocol for *M. polymorpha* thallus transformation, further enhancing the ability to introduce multiple genetic modifications and advance plant engineering.

In summary, in this study, we outline the generation and validation of constructs for gRNA multiplexing in *M. polymorpha*, providing a straightforward and efficient pipeline for multiplex gene editing.

## Results

### Synthesis of multiplex tRNA-gRNA modules using Type II S cloning

We adapted the protocol originally developed for rice (Xie, Minkenberg, and Yang 2015) to clone tRNA-gRNA modules using the Loop assembly Type II S cloning (Pollak et al. 2019; Sauret-Güeto et al. 2020) to enable efficient gene editing in *M. polymorpha* (**Figure 1 and 2**). The process begins with primer design and PCR amplification of the tRNA-gRNA parts from the pGTR plasmid (Xie et al. 2015; Addgene #63143) which contains the tRNA and gRNA sequences (**Figure 2A**). The PCR products contain the tRNA sequence, the gRNA scaffold and a part of the gRNA protospacer sequence. These PCR products are then fused together to reconstitute multiplexed tRNA-gRNA-protospacer-gRNA-scaffold modules (up to three in our trials but the systems allows to combine a greater number) and placed under the control of the Mp*U6* promoter in a L1 vector (OpenPlant plasmids OP-074, Addgene #136136 or OP-075, Addgene #136137 (Sauret-Güeto et al. 2020)) via Loop assembly cloning (**Figure 2B**). After transformation of the L1 Loop reaction into *E. coli* cells, screening 1-3 colonies was sufficient to identify correctly assembled constructs in our trials. Finally, the L1 tRNA-gRNA construct is transferred into the L2 pCsA (Addgene #136067 (Sauret-Güeto et al. 2020)) acceptor vector through another round of Loop assembly (**Figure 2C**), linking the tRNA-gRNA unit with the transcription unit for Cas9 expression (Addgene #136135 (Sauret-Güeto et al. 2020)) and a transcription unit containing the desired selection marker. In summary, this protocol enables simple, fast and efficient generation of constructs for targeted gene editing in *M. polymorpha*.

**Figure 2:**
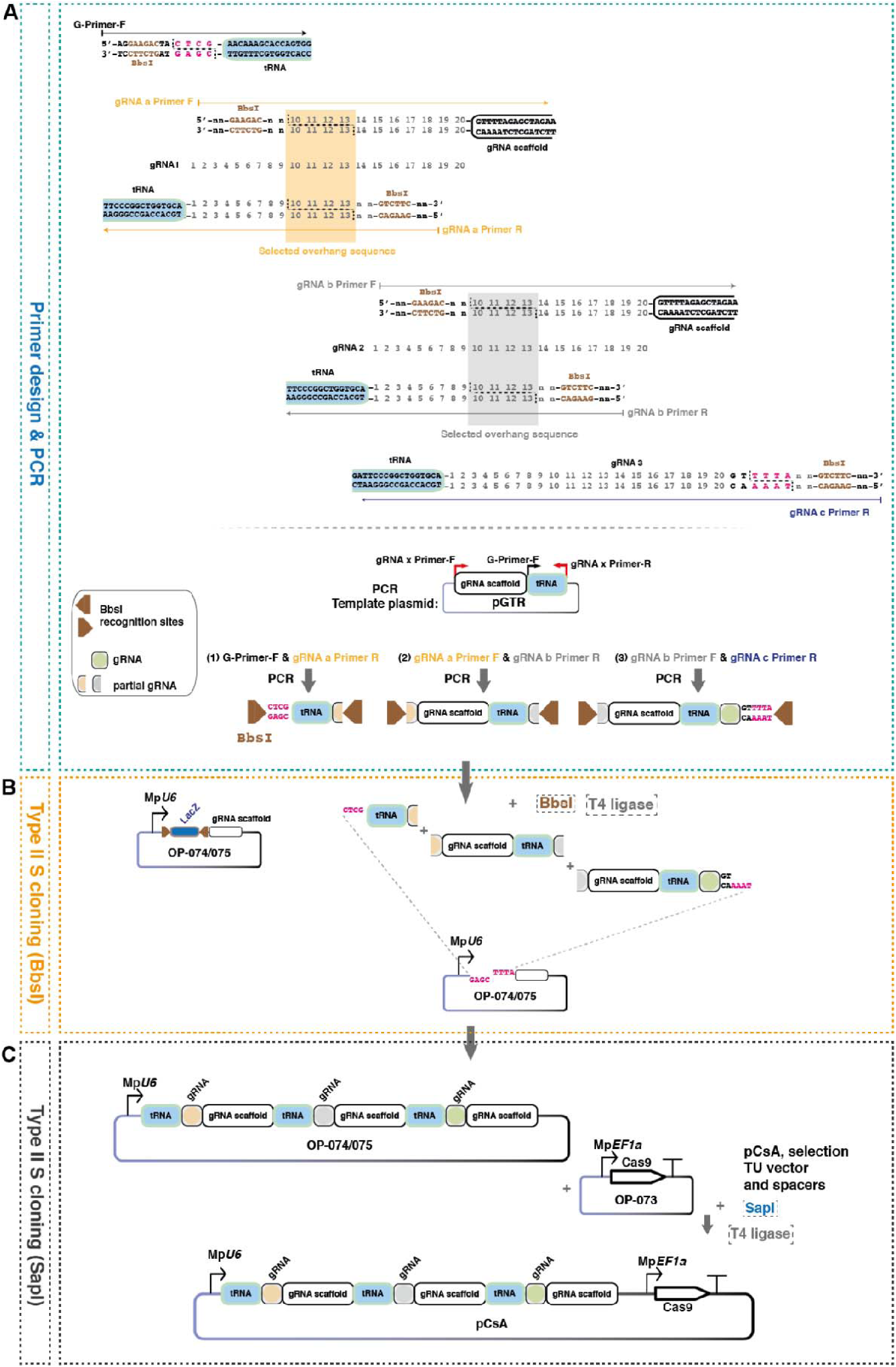
Schematic diagram of the workflow for generating a tRNA-mediated multiplex gRNA expression construct. **(A)** Top: The first primer includes a portion of the tRNA sequence along with a BbsI recognition site and four-base overhang sequences for Loop assembly. The middle primers contain BbsI recognition sites and four-base overlapping overhang sequences for Loop assembly, which can correspond to any four consecutive nucleotides (highlighted with colored rectangles) within the 20-nucleotide gRNA sequence (shown with numbers from 1 to 20). Forward middle primers also contain a portion of the gRNA scaffold sequence, whereas reverse middle primers contain a portion of the tRNA sequence. The final primer includes part of the tRNA sequence, the full reverse-complement of the last gRNA, and a BbsI recognition site plus four-base overhang sequences for Loop assembly. Overhangs for cloning into the acceptor vectors shown with pink letters. Bottom: The pGTR plasmid should be used as the template for the PCR reactions. The primer combinations used for amplification are as follows: (1) G-primer-F & gRNA a Primer R, (2) gRNA a Primer F & gRNA b Primer R, and (3) gRNA b Primer F & gRNA c Primer R, n: any nucleotide (please see **Supplemental Information** for a detailed explanation and an example of primer design). **(B)** After PCR amplification and gel extraction, all fragments are combined with the OP-074 or OP-075 vector (Sauret-Gueto et al., 2020) in a Loop Assembly/Type IIS cloning reaction (see **Supplemental Information** for a detailed protocol of Loop Assembly L1 cloning). LacZ: lacZα cassette for blue-white screening of colonies (negative blue colonies contain undigested L1 vectors, while positive white colonies contain tRNA-gRNA parts inserted into the L1 vectors. **(C)** Finally, the tRNA-gRNA-OP-074 or OP-075 vector is combined with the OP-073 vector, which contains the Mp*EF1a::Cas9* transcription unit, an appropriate L1 vector for plant selection, in a Loop Assembly/Type IIS cloning reaction (see **Supplemental Information** for a detailed protocol of Loop Assembly L2 cloning).

### tRNA-gRNA multiplexing improves genome editing efficiency

To test efficiency of the new system, we targeted the *M. polymorpha GOLDEN-2 LIKE* (Mp*GLK*) gene, mutations of which lead to reduced levels of chlorophyll (Yelina et al. 2024; Frangedakis et al. 2024; Hernández-Muñoz et al. 2024), and result in an easily observable, non-lethal phenotype which is ideal for the evaluation of CRISPR/Cas9 efficiency.

Using the tRNA-gRNA system, we constructed vectors to express one or three gRNAs to target the Mp*GLK* locus to generate large deletions (**Figure 3A**). Additionally, as a control, we included a vector allowing gRNA expression from the commonly used Mp*U6* promoter without tRNA sequences (**Figure 3A**). We transformed all three constructs into sporelings. In all cases, we observed regenerated thalli with pale green sectors, indicating successful disruption of the Mp*GLK* gene. We classified the resulting plants into three distinct phenotypic categories: wild-type, chimeric, and fully mutant (**Figure 3A**). For the control vector, expressing the gRNA without tRNA sequences, we observed approximately 88% wild-type, 10% chimeric, and 2% fully mutant plants (**Figure 3B**). These numbers were comparable to those obtained when expressing the same gRNA with the tRNA sequences (**Figure 3B**). Upon delivering the three gRNAs targeting Mp*GLK*, approximately 55% of the primary transformants exhibited wild-type, 33% chimeric, and 12% fully mutant phenotypes (**Figure 3B**). To confirm successful gene disruption, we performed Sanger sequencing, which validated the expected mutations (**Figure 3C-D and Supplemental Figure 1A)**. To estimate the frequency of large deletions, we genotyped 60 independent mutant lines and found that 10% harbored large deletions (∼700-800 bp, between gRNA3&7) (**Figure 3E and Supplemental Figure 1A)**. Overall, our results demonstrate a statistically significant increase in gene-editing efficiency when using tRNA-gRNA multiplexing compared to the use of vectors expressing a single gRNA, underscoring the effectiveness and reliability of our system.

**Figure 3:**
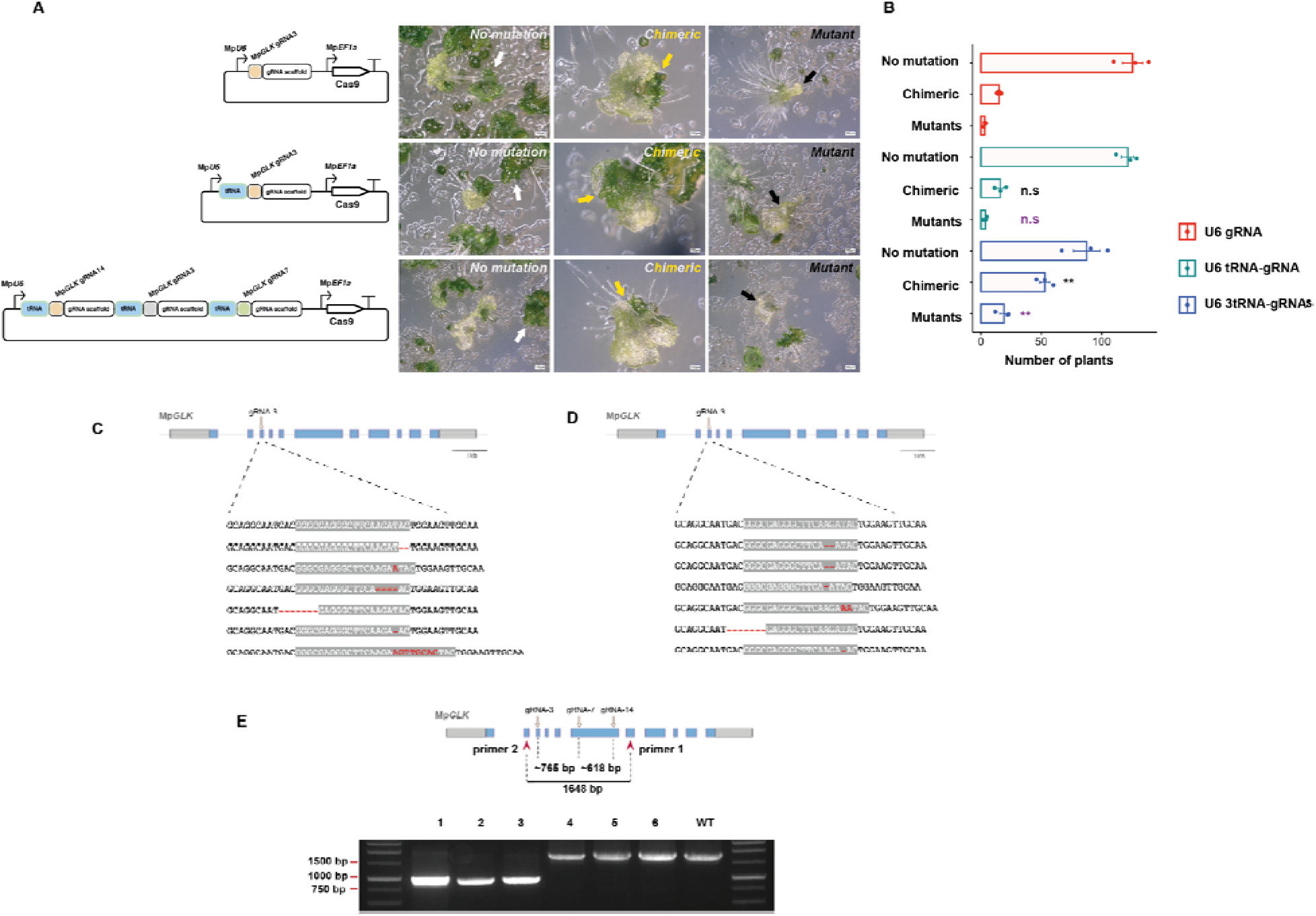
Evaluation of mutant generation using the tRNA-mediated multiplex gRNA expression method. **(A)** Left: Schematic representation of the plant transformation vector for simultaneous delivery of one or three gRNAs and Cas9 into *M. polymorpha* plants by *Agrobacterium*-mediated transformation, with or without the use of tRNAs. Right: The three phenotypic classes of regenerating *M. polymorpha* plants: wild-type, chimeric, and mutant. Scale bar: 400 μm. **(B)** Comparison of the number of transformants with wild-type, chimeric, or mutant phenotypes. Graphs show values from triplicate experiments (dots) and their average (bars). Error bars represent the SEM; n = 3. A pairwise t-test was conducted to compare the number of chimeric mutants (generated using either the U6 tRNA-gRNA vector or U6 tRNA-gRNA vector) against the number of plants generated using the control vector (U6 gRNA). ** for p < 0.01, while no significant difference (n.s) was noted for p > 0.05. A similar pairwise test was performed for the fully mutated plants. **(C-D)** Sequence analysis of Mp*glk* mutant lines. Schematic representation of Mp*GLK* gene structure showing exons as blue rectangles, untranslated regions (UTRs) as grey rectangles, and introns as grey lines. The position of the gRNA used for CRISPR/Cas9 gene editing is indicated with an arrow. The wild-type *M. polymorpha* Cam-1 sequence is shown at the top, with the 20-bp gRNA target sequence highlighted in grey. Mutations are highlighted in red. **(E)** Gel electrophoresis images of PCR-based genotyping of Mp*glk* mutants obtained using the vector that allows the expression of three gRNAs. The positions of primers used are shown with red arrowheads. The expected size of PCR products when large deletions occurred is approximately ∼600-800 bp (lanes 1-3). The expected size of PCR products for wild-type or mutants with small deletions is 1648 bp (lanes 4, 5, 6 and WT).

### A simplified and optimized thallus transformation method for *M. polymorpha*

Stable transformation of *M. polymorpha* can be obtained using either germinating spores or thallus fragments (Iwakawa et al. 2021; Kubota et al. 2013; Ishizaki et al. 2008). Spore-based transformation offers high efficiency but requires approximately three months for spore production and can be challenging for certain *M. polymorpha* accessions. Furthermore, transformants derived from spores consist of a mix of male and female plants and are not isogenic. In contrast, thallus transformation uses thallus fragments or gemmalings as tissue material, which are more easily obtained than the spores and yield an isogenic population of transformants. Thallus transformation also allows the introduction of additional transgenes into specific mutant strains without segregation, as it bypasses sexual reproduction.

Here, we present a simplified protocol for *M. polymorpha* thallus transformation, adapted from a method originally developed for hornwort thallus transformation (Waller et al. 2023). This protocol utilizes the 2-(N-morpholino)ethanesulfonic acid (MES) buffer to maintain a stable pH in the co-cultivation medium. Maintaining a stable pH of 5.5 during co-cultivation has been shown to significantly enhance *Agrobacterium*-mediated infection in *Arabidopsis thaliana* and also hornworts, likely by inhibiting calcium-mediated defense signaling (Y.-C. Wang et al. 2018; Waller et al. 2023). This optimized protocol omits the need for sucrose (Kubota et al. 2013) treatment of the regenerating tissue prior to co-cultivation with the *Agrobacterium* and reduces the required volumes of co-cultivation media, effectively miniaturizing the process.

The protocol consists of the following steps (**Figure 4A&B and Supplemental Figure 2A**): Three two-weeks-old gemmalings were transferred to a well of a 6-well plate, containing 5 mL of liquid KNOP medium supplemented with 1% (w/v) sucrose and 10mM MES, pH 5.5. The gemmalings were then fragmented with a scalpel (into approximately 20-25 fragments) and acetosyringone was added. The tissue was co-cultivated with *Agrobacterium tumefaciens* carrying an *eGFP* gene fused to a cytoplasm targeting signal and hygromycin selectable marker gene, for three days on a shaker at 110 rpm, under ambient light and 21°C. After co-cultivation, thallus fragments were transferred to growth media containing hygromycin or the selectable marker of choice. Successful transformants, identifiable by eGFP fluorescence under a dissecting microscope, became visible from three to six weeks (**Figure 4C-G**).

**Figure 4:**
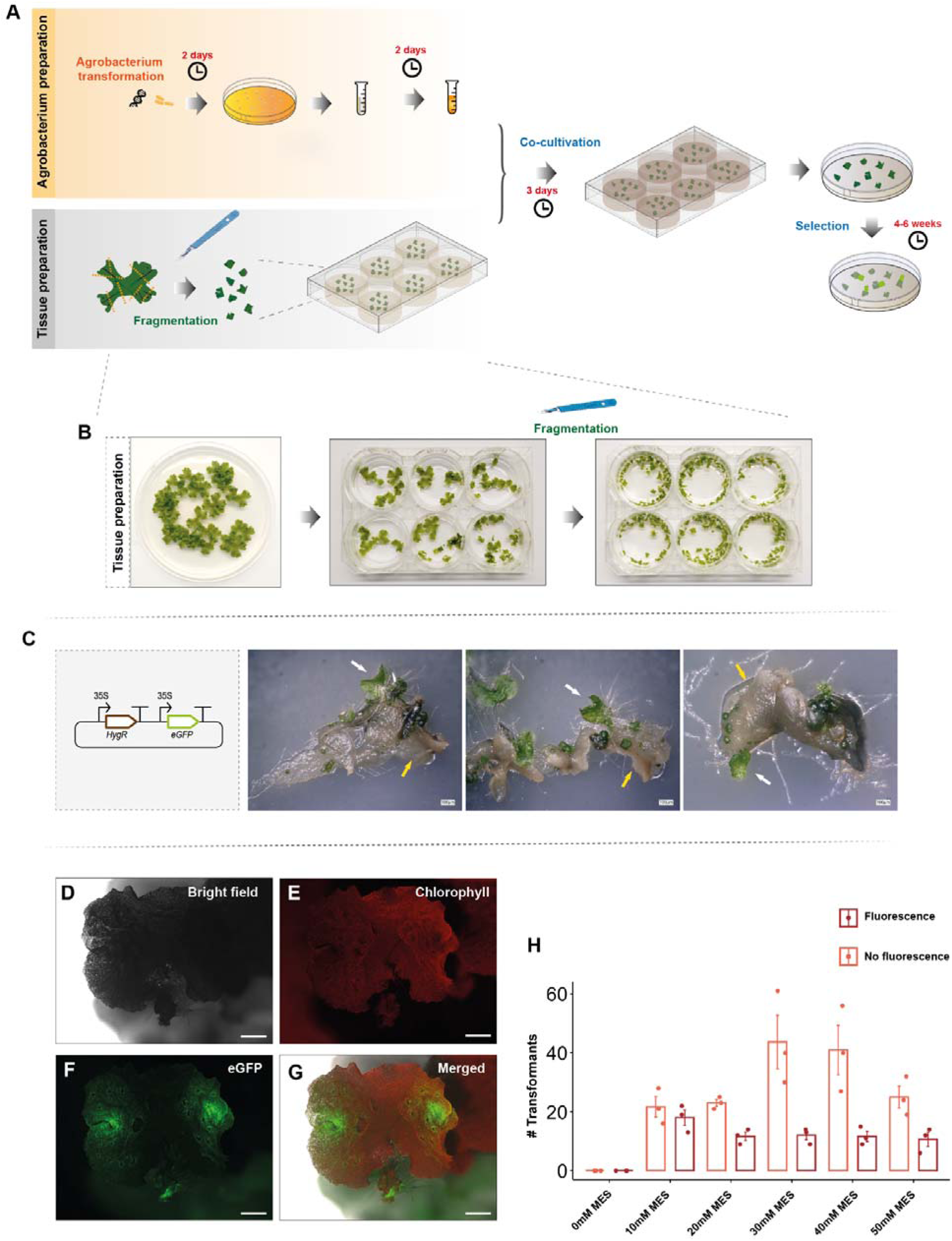
Optimized *M. polymorpha* thallus transformation protocol. **(A)** Outline of the thallus transformation protocol steps: Thallus tissue is transferred into a 6-well plate containing 5 mL of co-cultivation medium and then fragmented with the aid of a sterile scalpel. The tissue is co-cultivated with *Agrobacterium* for three days and then spread on antibiotic-containing growth medium. After approximately 4–6 weeks, putative transformants are visible as regenerating green tissue. **(B)** Representative images of *M. polymorpha* 2-week-old gemmae, which are then transferred into a 6-well plate and fragmented prior to the addition of *Agrobacterium*. **(C)** Schematic representation of constructs for the expression of two transcription units (TUs): one TU for the expression of the *hygromycin B phosphotransferase* (*hph*) gene under the control of the cauliflower mosaic virus (CaMV) 35S promoter, and one TU for the expression of eGFP under the control of the CaMV 35S promoter, targeted to the plasma membrane using the Lti6b localization signal. Representative images of *M. polymorpha* thallus fragments after 4 weeks on antibiotic-containing growth medium show regenerating tissue (white arrows) that is potentially successfully transformed. Untransformed, dying tissue is indicated with yellow arrows. Scale bar: 100 μm. **(D-G)** Images of *M. polymorpha* thallus tissue expressing p-35S::eGFP-Lti6b for plasma membrane localization. Scale bar: 1 mm. **(H)** Comparison of the number of transformants (per 20 thallus fragments) under different MES concentrations in the co-cultivation medium (initial pH set to 5.5). Graphs show values from triplicate experiments (dots) and their average (bars). Error bars represent the SEM; n = 3.

To further optimize the protocol, we evaluated the effect of MES concentration on transformation efficiency by adding it to the co-cultivation medium at concentrations ranging from 0 to 50 mM. eGFP fluorescence was used to estimate transformation efficiency (**Figure 4C**). Increasing MES concentration to 30 mM resulted in more transformants, although concentrations above 40 mM led to a decrease in the total number of recovered transformants. The optimal MES concentration was therefore determined to be 30 mM. Interestingly, in many cases a single thallus fragment gave rise to more than one regenerated fragment/putative transformant (**Figure 4C**). However not all regenerated fragments showed fluorescence potentially due to a partial T-DNA transfer or gene silencing (**Figure 4D-H**). On average, we observed a transformation efficiency of approximately 30 transgenic plants per 20 fragments across different batches (**Figure 4H**). All subsequent trials were performed using a MES concentration of 30 mM. The new method can be applied not only to gemmaling tissue but also to more mature thallus tissue, increasing our method’s flexibility and ease of use. However, under our conditions, efficiency was reduced in older tissue (**Supplemental Figure 2B**). To address the reduced efficiency, scaling up the protocol using co-cultivation in flasks could be an alternative. Overall, our updated thallus transformation method facilitates the straight-forward generation of isogenic transformants, offering a rapid approach for genetic studies and functional analyses in *M. polymorpha*.

### The optimized thallus transformation allows efficient generation of single and double mutants

Next, we tested whether our improved thallus transformation system could be used to obtain CRISPR/Cas9 mutants. To do this, we used the same vector as was used for the sporeling transformation trials to express three gRNAs targeting the Mp*GLK* gene (**Figure 5A**). Similarly to sporeling transformation, the plants resulting from thallus transformation fell into three distinct phenotypic categories: wild-type, chimeric, and fully mutant (**Figure 5B**). We transformed approximately 200 thallus fragments and 74%, 23% and 3% of the primary transformants (n=100) exhibited wild-type, chimeric, and fully mutant phenotypes respectively (**Figure 5C**). Notably, chimeric plants displayed multiple sectors with the mutant phenotype, indicating that mutations in each sector arose independently (**Figure 5B).** Mutant sectors mostly appeared close to the meristematic notch **(Supplemental Figure 3A),** possibly due to the stronger activity of the Mp*Ef1a* promoter that drives Cas9 expression in our constructs (Tse et al. 2024; Althoff et al. 2014). However, we also noticed that mutant sectors can arise in more mature areas of the thallus **(Supplemental Figure 3B)**.

**Figure 5:**
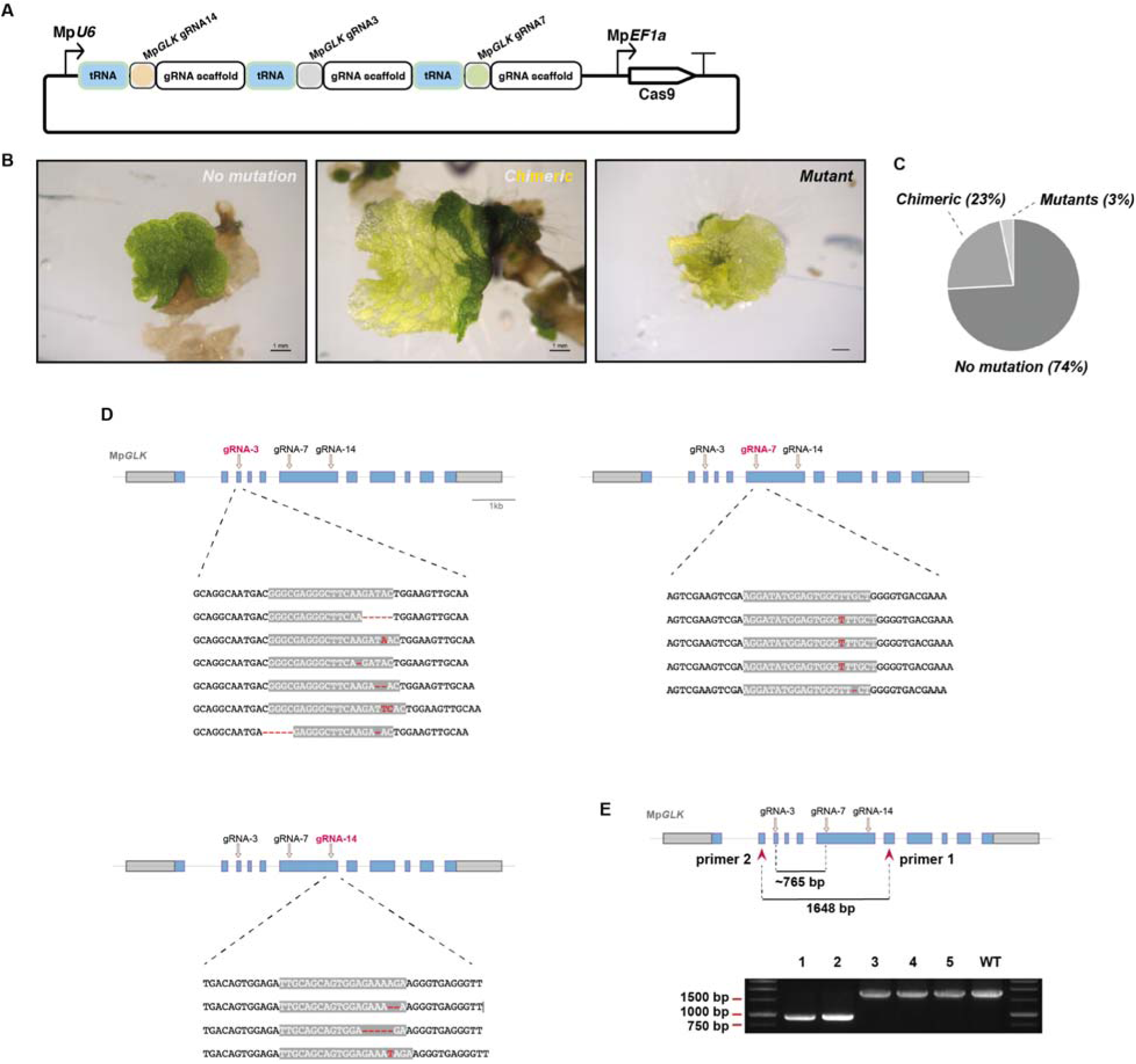
Generation of mutants using the optimized thallus transformation. **(A)** Schematic representation of the plant transformation vector for simultaneous delivery of three gRNAs and Cas9 into *M. polymorpha* plants by *Agrobacterium*-mediated transformation, with or without the use of tRNAs. **(B)** The three phenotypic classes of regenerating *M. polymorpha* plants: wild-type, chimeric, and mutant. Scale bar: 1 mm. **(C)** Comparison of the number of transformants with wild-type, chimeric, or mutant phenotypes. n = 100. **(D)** Sequence analysis of Mp*glk* mutant lines. Top: Schematic representation of Mp*GLK* gene structure. The position of the gRNA used for CRISPR/Cas9 gene editing is shown with an arrow. Bottom: Mutant genotyping analysis. The wild-type *M. polymorpha* Cam-1 sequence is shown at the top, with the 20-bp gRNA target sequence highlighted in grey. Mutations are highlighted in red. **(E)** Gel electrophoresis images of PCR-based genotyping of Mp*glk* mutants obtained using the vector that allows the expression of three gRNAs. Schematic representation of Mp*GLK* gene structure showing exons as blue rectangles, untranslated regions (UTRs) as grey rectangles, and introns as grey lines. Positions of primers used are shown with red arrowheads. The expected size of PCR products when large deletions occur is approximately 765 bp (lanes 1 and 2). The expected size of PCR products for wild-type or mutants with small deletions is 1648 bp (lanes 3, 4, 5 and WT).

To verify successful gene disruption, we conducted Sanger sequencing, which confirmed the expected mutations (**Figure 5D-E and Supplemental Figure 1B)**. To evaluate the occurrence of large deletions, we genotyped 60 independent mutant lines and observed that 2% carried a ∼765 bp deletion (gRNA3&7) (**Figure 5E**), which was confirmed with Sanger sequencing **(Supplemental Figure 1B).**

We also assessed whether our improved thallus transformation method could be used to generate double mutants. In this approach, thalli from a mutant plant were subject to secondary transformation using an L2 construct containing a different selectable marker and a transcription unit driving expression of a gRNA targeting a second gene (in this case Mp*RR-MYB5*) (**Figure 6A**). There was no need to include a separate transcription unit for Cas9 expression, as it was already present in the mutant background, simplifying the process. We selected the Mp*RR- MYB5* gene as a target, because, similarly to Mp*GLK*, its mutations lead to reduced chlorophyll, and, therefore, an easily observable, non-lethal phenotype ideal for the evaluation of gene editing efficiency(Frangedakis et al. 2024). We targeted Mp*RR-MYB5* in two different mutant backgrounds, Mp*gata4* and Mp*glk* (Yelina et al. 2024) (**Figure 6B&C)**. We used two different mutant backgrounds with distinct thallus morphologies and phenotypes to test the effectiveness of our protocol across varying mutant thallus types. The resulting plants again fell into three distinct phenotypic categories: original mutant background, chimeric, and fully mutant (**Figure 6B&C)**. We transformed approximately 200 thallus fragments of Mp*gata4* or Mp*glk* mutant plants. Of the recovered transformants (n=100), 85% were wild type for Mp*RR-MYB5*; 12% were chimeric and 3% were fully mutant for Mp*rr-myb5* (**Figure 6D).** Out of the transformants recovered from transforming Mp*glk* mutant thallus fragments (n=100), 78% were wild type for *MpRR-MYB5*; 20% were chimeric and 2% were fully mutant for Mp*rr-myb5* (**Figure 6E**). Successful gene disruption was confirmed by Sanger sequencing (**Figure 6D&E)**. Collectively, our results demonstrate that the optimized thallus transformation system is a reliable and efficient method for generating both single and double CRISPR/Cas9-mediated mutants, including in mutant backgrounds with altered thallus tissue morphology or development.

**Figure 6:**
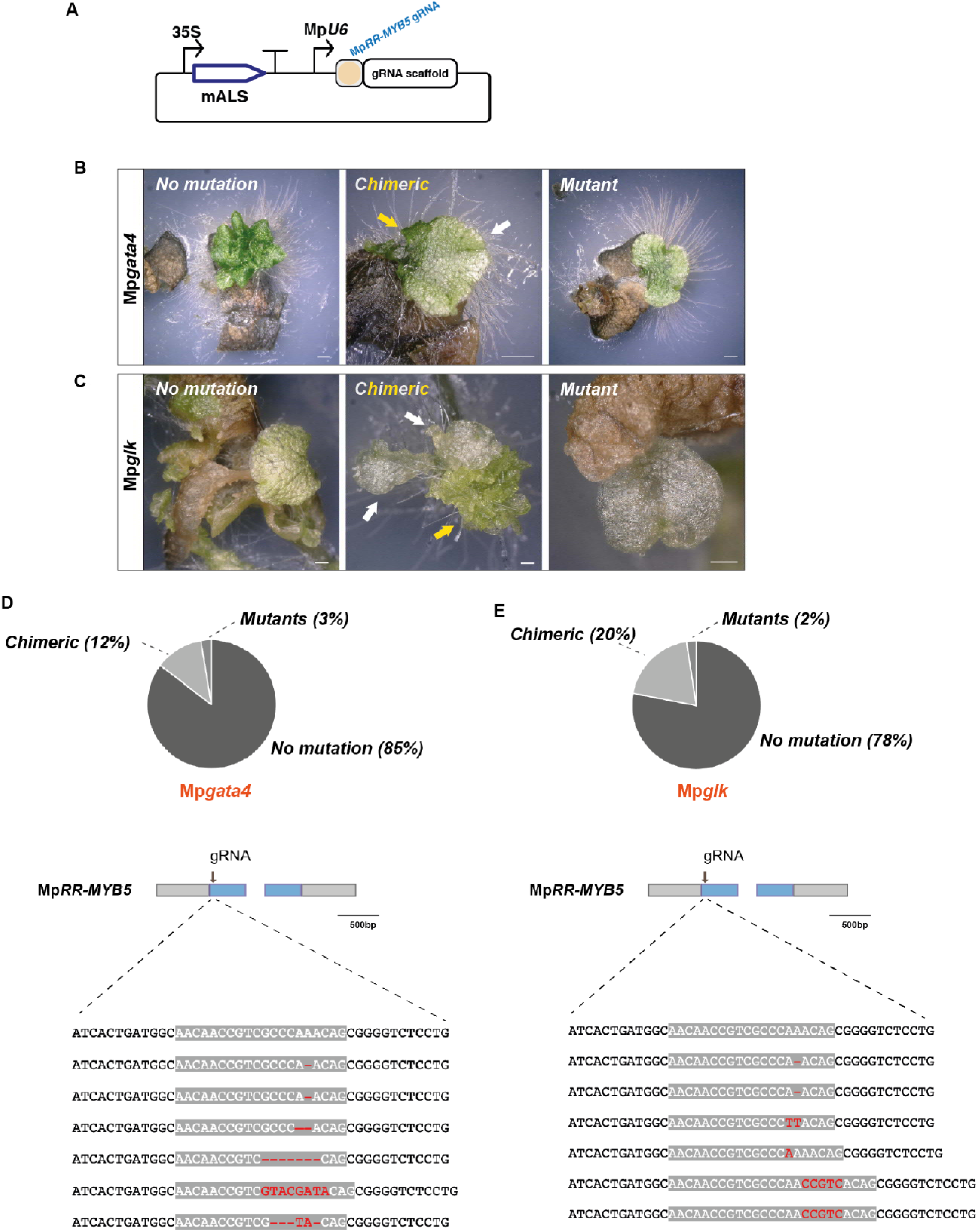
Generation of double mutants using the optimised thallus transformation. **(A)** Schematic representation of the plant transformation vector for delivery of a gRNA for targeting Mp*RR-MYB5* into Mp*gata4* or Mp*glk* mutant *M. polymorpha* plants by *Agrobacterium*- mediated transformation. mALS: Chlorsulfuron selection. **(B-C)** The three phenotypic classes of regenerating Mp*gata4* and Mp*glk* mutant plants transformed with the L2 vector (A) for targeting Mp*RR-MYB5*: Mp*gata4*, chimeric, and mutant or Mp*glk*, chimeric, and mutant. Scale bar: 1 mm **(D-E)** Top: Schematic representation of Mp*RR-MYB5* gene structure showing exons as blue rectangles, untranslated regions (UTRs) as grey rectangles and introns as grey lines. Positions of gRNAs used for CRISPR/Cas9 gene editing are shown as arrows. Middle: Comparison of the number of transformants with Mp*glk,rr-myb5* or Mp*gata4*,*rr-myb5* chimeric or mutant phenotypes. n=100. Bottom: Sequence analysis of Mp*glk,rr-myb5* and Mp*gata4*,*rr-myb5* double mutant lines. The wild-type *M. polymorpha* Cam-1 sequence is shown at the top, with the 20 bp gRNA target sequence highlighted with grey. Mutations are highlighted in red.

## Conclusions

CRISPR-based genome editing tools have significantly advanced biology (Chen et al. 2019; J. Y. Wang and Doudna 2023; Pacesa, Pelea, and Jinek 2024), and their application to *M. polymorpha* (Sugano et al. 2018; Sauret-Güeto et al. 2020; Sugano et al. 2014) has enhanced substantially this model organism’s utility.

In this study, we successfully adapted gRNA multiplexing by expressing gRNAs from a single transcriptional tRNA-gRNA unit (Xie, Minkenberg, and Yang 2015) for CRISPR/Cas9 genome editing in *M. polymorpha*. By incorporating tRNA sequences into the OpenPlant CRISPR/Ca9 kit vector system, we demonstrated that tRNA-gRNA approach results in functional gRNAs. This approach streamlines genome editing in *M. polymorpha*, allowing the introduction of multiple modifications in a single transformation event. We validated the effectiveness of this strategy through successful targeted mutagenesis, including large deletions readily detectable by PCR, a feature that simplifies mutant screening. Finally, our method can also facilitate the simultaneous targeting of multiple genes or can be adapted to other *M. polymorpha* CRISPR/Cas9 vector systems.

Beyond fundamental research, *M. polymorpha* is a promising chassis for synthetic biology applications (Sauret-Güeto et al. 2020; Frangedakis et al. 2021; Boehm et al. 2017). It offers several advantages, including a relatively fast life cycle, simpler tissue organisation compared to other plant models in addition to well characterized nuclear and plastid genomes which are both amenable to genetic manipulation and high-throughput approaches (Annese et al. 2025). To date, up to five different selectable markers have been tested in *M. polymorpha*, further expanding the capacity to introduce multiple transgenes (Ishizaki et al. 2015; Robinson et al. 2024). All these can facilitate the reprogramming of gene regulatory networks with high precision and speed. Moreover, *M. polymorpha* can potentially serve as a platform for testing and developing novel biotechnological tools, such as biosensors, bio-based production of high value compounds, and cell type specific metabolic engineering (Tansley, Patron, and Guiziou 2024; Yang and Reyna-Llorens 2023). Its ability to produce complex secondary metabolites further enhances its potential for bioengineering (Lindström Battle and Sweetlove 2025; Forestier et al. 2025). As a result, developing improved CRISPR/Cas9 tools in *M. polymorpha* can enable more versatile genome engineering.

In conclusion, our work establishes a new efficient approach for genome editing in *M. polymorpha* that uses the tRNA-gRNA system for simultaneous editing of multiple genes. This system represents a significant advancement in the genetic toolkit for *M. polymorpha*, with implications for plant development studies, evolutionary biology, and synthetic biology applications.

## Material and methods

### Plant material and growth conditions

*Marchantia polymorpha* accessions Cam-1 (male) and Cam-2 (female) (Delmans, Pollak, and Haseloff 2017) were used in this study. Plants were grown and maintained on ½ strength Gamborg’s B5 medium plus vitamins (DUCHEFA, #G0210.0050) and 1.2 % (w/v) agar (MELFORD, #A20021), under continuous light at 21°C.

### *M*. *polymorpha* axenic cultures for spore production

*M. polymorpha* thallus fragments or gemmae were transferred into autoclaved microboxes containing 15 “Jiffy 7” pellets (JIFFY PRODUCTS INTERNATIONAL BV), as previously described (Sauret-Güeto et al. 2020). Briefly, 200 mL of water was added, and the plants were allowed to grow for one month under continuous light at 21°C. After this period, far-red light was applied to induce the development of reproductive organs. Within 3–4 weeks, mature gametophores, ready for fertilization, became visible. For fertilization, a drop of sterile water was added to the male gametophore, and the released sperm was transferred with a pipette onto the female gametophore. After an additional month, mature sporophytes were visible and ready for collection.

### Single gRNA cloning into L1 vectors

Two oligos (primers) for cloning the gRNA target sequence were designed, as previously described (Sauret-Güeto et al. 2020). To prepare the annealing reaction, 1 μL of each oligo (100 μM) was mixed with 8 μL of water to a final volume of 10 μL. The mixture was then annealed in a thermocycler under the following conditions: 37 °C for 30 minutes, 95 °C for 5 minutes, followed by a gradual decrease to 25 °C at a rate of 5 °C per minute. After annealing, the gRNA target sequence was cloned into the L1_lacZgRNA-Ck2 or L1_lacZgRNA-Ck3 vectors. L1 BbsI-mediated Loop reactions were performed with minor modifications to previous protocols (Sauret-Güeto et al. 2020). Briefly, the reaction included 7.5 nM (1 μL) of pCk vectors and 2 μL of annealed oligos. The Level 1 Loop assembly master mix contained 2 μL of 10× T4 DNA ligase buffer (NEB), 1.5 μL of 1 mg/mL bovine serum albumin (NEB, #B9200S), 1.5 μL of T4 DNA ligase 400 U/μL (NEB, #M0202S), 1.5 μL of BbsI-HF 10 U/μL (NEB, #R3539), and nuclease-free HLO to a final volume of 20 μL. The cycling conditions included 26 cycles of 37 °C for 3 minutes and 16 °C for 4 minutes, followed by enzyme inactivation at 50 °C for 5 minutes and 80 °C for 10 minutes. For transformation, 20 μL of chemically competent *E. coli* cells were incubated with 10 μL of the Loop assembly reaction and plated on LB agar containing 50 μg/mL kanamycin (MELFORD, #K22000–1.0) and 40 μg/mL X-Gal (5-bromo-4-chloro-3- indolyl-β-D-galactoside) (ThermoFisher, #15520018).

### Primer design for tRNA-gRNA module amplification

The tRNA-gRNA spacer-specific primers were designed with BbsI 4 bp overlapping overhangs for Golden Gate/Loop Assembly. The sequences for these primers are as follows (**Figure 2A**): G-primer-F (AGgaagacTACTCGAACAAAGCACCAGTGG) and “gRNA c Primer R” (GAgaagacTATAAAAC-LAST-gRNA-REVcompl-TGCACCAGCCGGGAATC). The sequences for “gRNA a Primer F” and “gRNA b Primer F” are GAgaagacAT-partial-gRNA- GTTTTAGAGCTAGAA, while “gRNA a Primer R” and “gRNA b Primer R” are GAgaagacTA- partial-gRNA-REVcompl-TGCACCAGCCGGGAA. The primer combinations used for amplification (when three gRNAs are combined, but can adapted to combine a larger number) are as follows: (1) G-primer-F with gRNA a Primer R, (2) gRNA a Primer F with gRNA b Primer R, and (3) gRNA b Primer F with gRNA c Primer R. The pGTR plasmid should be used as the template for these reactions. For detailed instructions and an example of primer design please see **Supplemental Information**. For PCR amplification Phusion polymerase (ThermoFisher, #F534S) was used according to the manufacturer’s instructions. PCR products were gel extracted using the QIAquick Gel Extraction Kit (QIAGEN, #28704).

### tRNA-gRNA cloning into L1 vectors

To clone the tRNA-gRNA DNA fragments into either L1_lacZgRNA-Ck2 or L1_lacZgRNA-Ck3, the same procedure used for single gRNA cloning was followed, with the only modification being the replacement of the annealed oligos with gel-extracted fragments. Reaction volumes were adjusted accordingly (see **Supplemental Information** for detailed protocols).

### L2 SapI mediated Loop Assembly

For the SapI-mediated Type IIS assembly, the following reaction was prepared: 2 μL of 10× T4 DNA ligase buffer (NEB), 1.5 μL of T4 DNA Ligase at 400 U/μL (NEB, #M0202S), 1.5 μL of SapI (10 U/μL, NEB, #R0569S), 1 μL (7.5 nM) of pCsA vector, 1 μL (15 nM) of L1 vectors, and nuclease-free H O to a final volume of 20 μL (please see Supplemental information for detailed instructions). After assembly, 10 μL of the reaction mixture was transformed into *E. coli* and plated on LB agar containing 100 μg/mL spectinomycin (MERCK, #S4014-5G) and 40 μg/mL X-Gal for selection.

### *Agrobacterium* culture preparation

Five milliliters of LB medium supplemented with rifampicin 50 μg/mL (DUCHEFA, #R0146), gentamicin 25 μg/mL (DUCHEFA, #G0124), and spectinomycin 100 μg/mL (Bio Basic, #SB0901), were inoculated with 3–4 *Agrobacterium tumefaciens* colonies (GV3101). The preculture was incubated at 28°C for 2 days at 100 rpm. After incubation (without measuring OD), 5 mL of the 2-day *Agrobacterium* culture was centrifuged at 2000 × g for 7 minutes. The supernatant was removed, and the pellet was resuspended in 5 mL of KNOP co-cultivation medium (0.25 g/ L KH_2_PO_4_, 0.25 g/L KCl, 0.25 g/L MgSO_4_•7H_2_O, 1 g/L Ca(NO_3_)2•4H_2_O and 12.5 mg/L FeSO_4_•7H_2_O) containing 1% (w/v) sucrose (MELFORD, #S24060-1000.0) and 100 μM acetosyringone (4′-Hydroxy-3′,5′-dimethoxyacetophenone) (MECK, #D134406). The culture was then incubated with shaking at 120 rpm at 28°C for an additional 3–5 hours.

### Transformation Procedure

Approximately 20 thallus cuttings, each around 2 × 2 mm - 3 × 3 mm, were transferred into a 6- well plate containing 5 mL of liquid KNOP co-cultivation medium supplemented with 1% (w/v) sucrose, 30 mM MES (2-(N-morpholino)ethanesulfonic acid) (pH 5.5), 80 μL of *Agrobacterium* culture, and acetosyringone at a final concentration of 100 μM. The tissue was co-cultivated with *Agrobacterium* on a shaker at 110 rpm at 21–22°C under ambient light for 3 days. After co-cultivation, the liquid was removed from the wells using a sterile plastic pipette, and the cuttings were transferred onto growth medium plates containing the appropriate antibiotic (hygromycin (MELFORD, #H7502) or chlorsulfuron (SIGMA, #34322)). To facilitate even distribution of thallus fragments on the plate, 2 mL of sterile water was added to each petri dish. After 3–6 weeks, successful transformation events were observed as small green regenerating tissue on dying or dead thallus fragments.

### Genotyping

For genotyping, small pieces (3 × 3 mm) of thalli from individual plants were placed in 1.5 mL Eppendorf tubes and crushed with an autoclaved micropestle in 80 μL of genotyping buffer (100 mM Tris-HCl, 1 M KCl, and 10 mM EDTA, pH 9.5). The tubes were incubated at 70°C for 25 minutes, followed by the addition of 350 μL of sterile water. A 2.5 μL aliquot of the extract was used as a template for PCR (using Phusion polymerase (ThermoFisher, #F534S) or Quick Taq ^®^ HS DyeMix (TOYOBO, #DTM-101)following the manufacturer’s instructions), and products were analyzed on a 1.5% (w/v) agarose gel. Primers used for genotyping: GLK_F: gaacctgcgtcgtgattgtag, GLK_R: gagttgttcgactgcctggac. For Mprr-myb5 genotyping we used the primers described in Frangedakis et al., 2024.

### Fluorescent Microscopy

A stereo microscope with fluorescence (Leica M205 FA) was used. The Leica M205FA was equipped with the following filters for observation of fluorescence: ET GFP (ET470/40 nm, ET525/50 nm), ET Chlorophyll LP (ET480/40 nm, ET610 nm LP), and ET GFP LP500 (ET470/40 nm, ET500 nm LP), used respectively for observing eGFP, chlorophyll, and eGFP together with chlorophyll autofluorescence.

### Light microscopy

Images were captured using a KEYENCE VHX-S550E microscope (VHX-J20T lens).

## Author contributions

E.F. and N.Y.E. designed the work. E.F., N.Y.E., S.K.E., F.R., and A.F., carried out the work.

J.H and J.M.H oversaw the project. E.F. wrote the manuscript with input from all authors.

## Acknowledgements

This work was funded as part of the BBSRC/EPSRC OpenPlant Synthetic Biology Research Centre Grant BB/ L014130/1 to J.H., BBSRC BB/F011458/1 for confocal microscopy to J.H, BBP0031171 to J.M.H. We thank Keiko Sakakibara from Rikkyo University, Japan, for the support during the last stage of the project.

## Notes

### Competing Interest Statement

The authors have declared no competing interest.

