## Supplemental Info for "A tRNA-gRNA multiplexing system for CRISPR genome editing in *Marchantia polymorpha*"

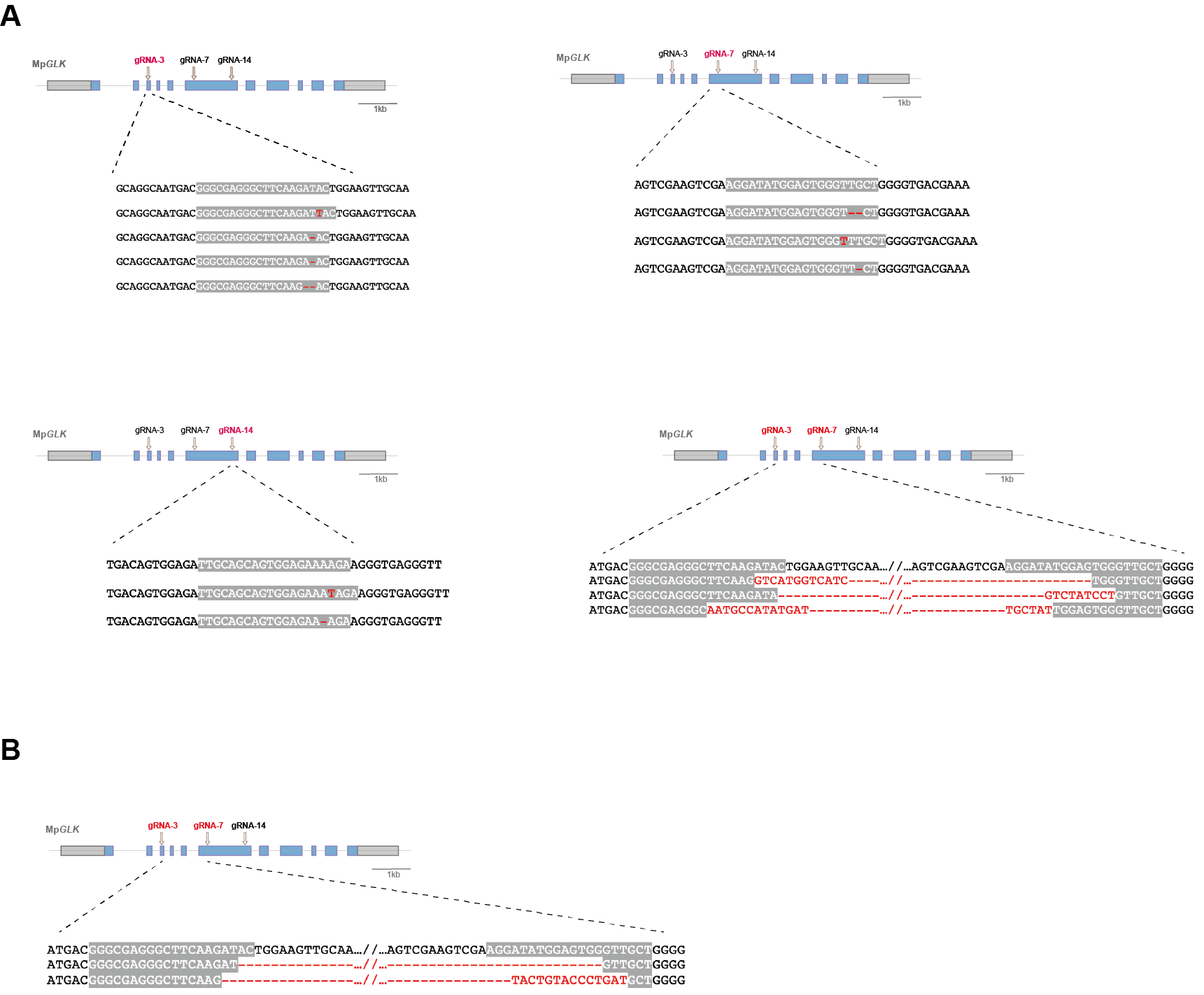

**Supplemental Figure 1:**

Sequence analysis of Mp*glk* mutant lines generated by transforming sporellings (A) or thallus fragments (B). Schematic representation of Mp*GLK* gene structure showing exons as blue rectangles, untranslated regions (UTRs) as grey rectangles and introns as grey lines. Position of gRNA used for CRISPR/Cas9 gene editing is shown with an arrow. The wild-type *M. polymorpha* Cam-1 sequence is shown at the top, with the 20 bp gRNA target sequence highlighted with grey. Mutations are shown with red.

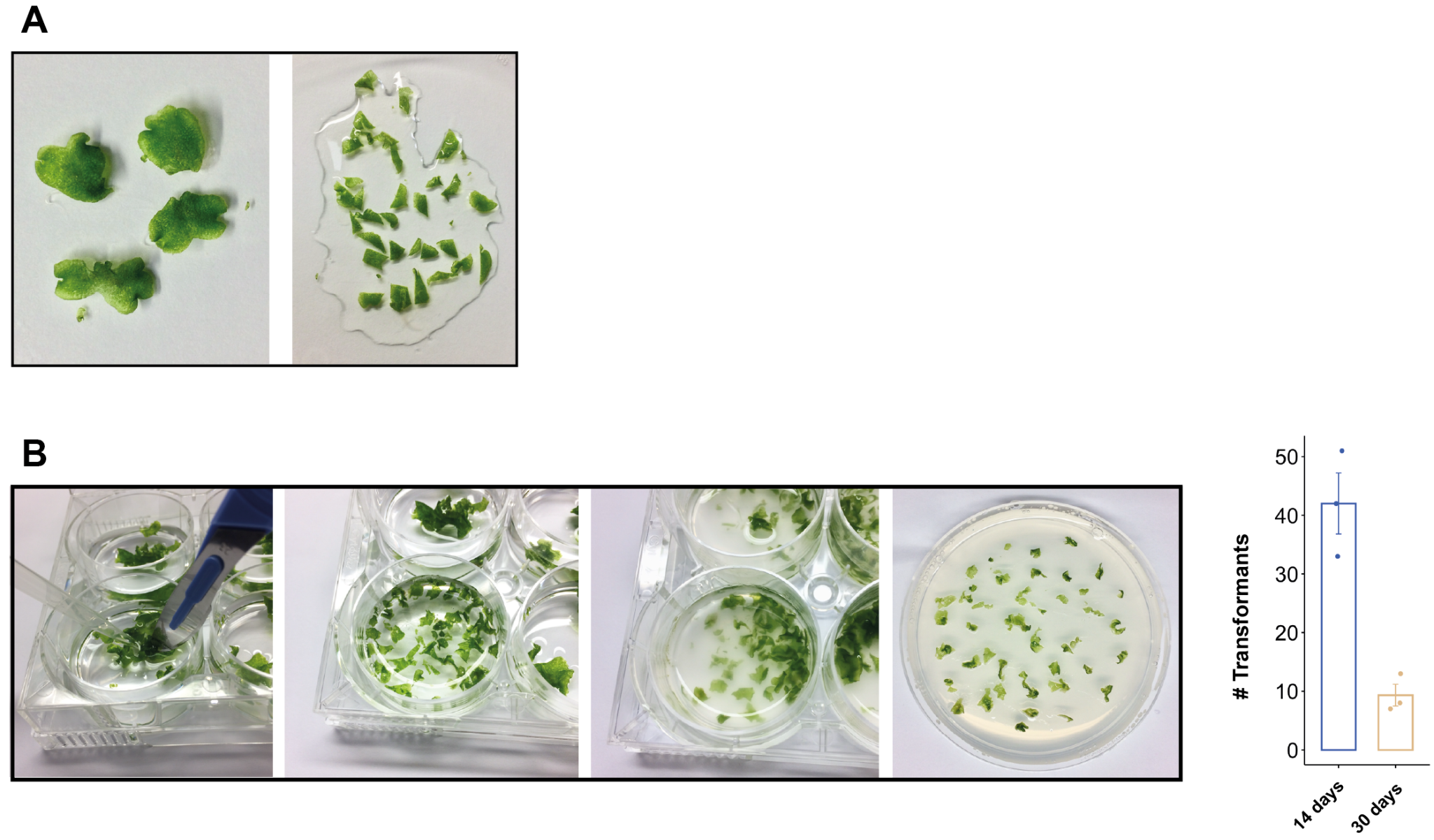

**Supplemental Figure 2:**

A) Representative images of *M. polymorpha* 2-week old gemallings, prior and after fragmentation

B) Left: Representative images of *M. polymorpha* 4-week old gemallings, that are then transferred into a six-well plate and fragmented prior to addition of the *Agrobacterium*. Right: Comparison of the number of transformants (per 20 thallus fragments) when using 14 days old or 30 days old gemmalings. Graphs show values of triplicate experiments (symbols) and their average (bars). Error bars values depicting the SEM; *n* = 3.

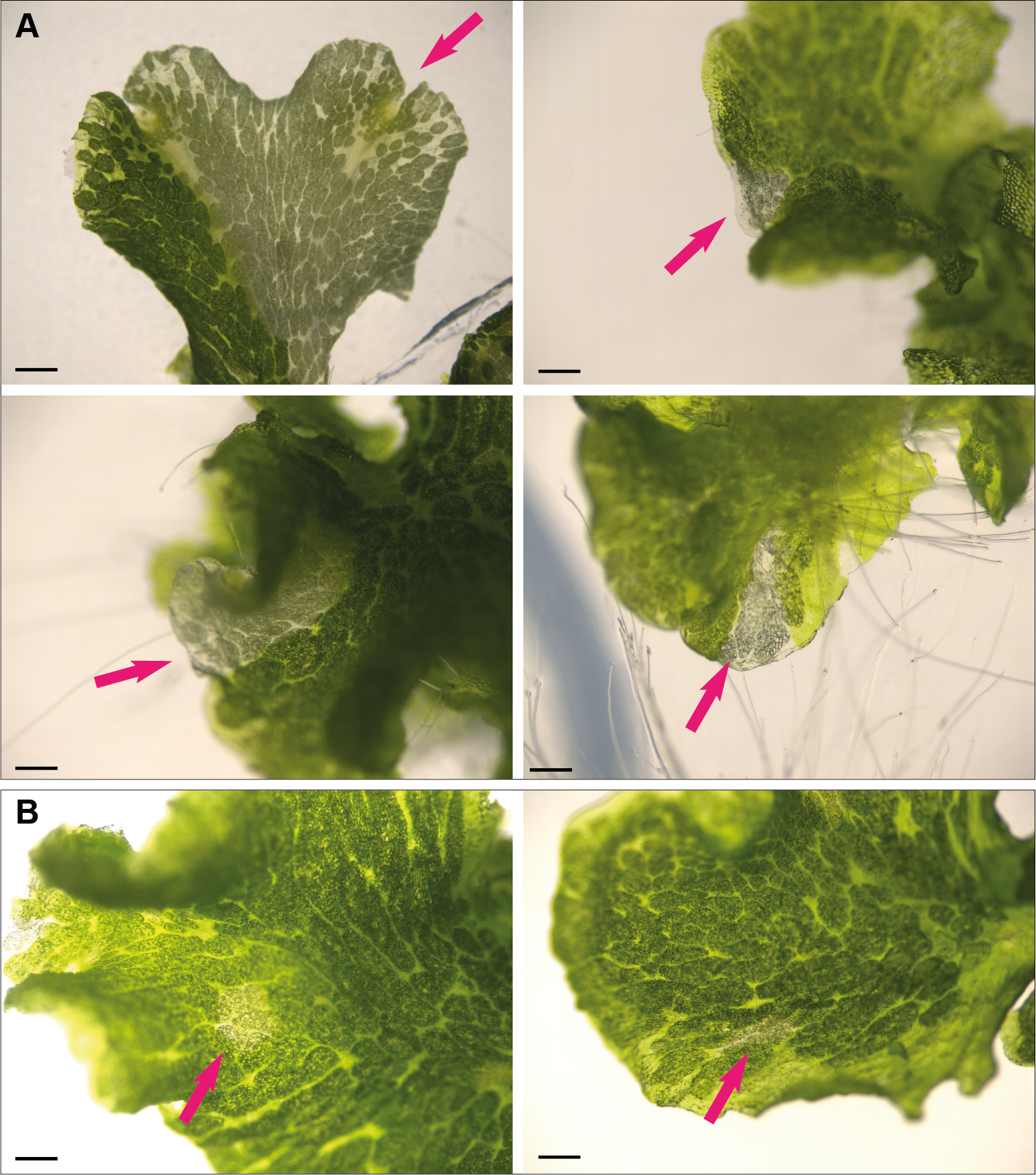

**Supplemental Figure 3:**

A-B) Examples of chimeric primary transformants with mutant sectors (arrows) in close proximity to the notch (A) and within the thallus (B). Scale bars: 1 mm

**Multiplex tRNA-gRNA constructs for *Marchantia polymorpha* CRISPR - AN EXAMPLE**

This section describes a *Marchantia polymorpha* specific modification of the Xie et al 2015 protocol (https://doi.org/10.1073/pnas.1420294112) for the synthesis of multicomplex tRNA-gRNA modules by Golden Gate Assembly/Loop Assembly. It has 3 main steps (see Main Figure 2 for a summary):

**A.** Primer design and PCR amplification of tRNA-gRNA fragments using the pGTR plasmid as the template.

**B.** Loop assembly cloning of the amplified tRNA-gRNA fragments into an L1 vector, to combine them with the Mp*U6* promoter (using plasmid OP-074 or OP-075).

**C.** Loop assembly cloning into the L2 pCsA acceptor vector to combine the *MpU6*::tRNA-gRNA unit with the transcription unit for Cas9 expression (from plasmid OP-073).

**A. Primer design and amplification of tRNA-gRNA parts using the pGTR plasmid as template**

The tRNA-gRNA spacer specific primers with 4 bp overlapping overhangs for BbsI Golden Gate/Loop Assembly should have the following sequences:

BbsI enzyme recognition sequence highlighted with orange

gRNA sequence shown with blue letters

Overhang sequences for cloning into the acceptor vector, highlighted with red

Sequence that is part of the gRNA scaffold or the tRNA highlighted with grey

G-primer-F

5’ AGgaagacTACTCGAACAAAGCACCAGTGG 3’

“gRNA c Primer R” in the main Figure 1:

5’ GAgaagacTATAAAAC-LAST-gRNA-REVcompl-TGCACCAGCCGGGAATC 3’

“gRNA a Primer F” and “gRNA b Primer F” in the main Figure 1:

5’ GAgaagacATxxxxxxxxxGTTTTAGAGCTAGAA 3’

“gRNA a Primer R” and “gRNA b Primer R” in the main Figure 1:

5’ GAgaagacTAxxxxxxxxxTGCACCAGCCGGGAA 3’

Primer combination is:

1 - G-primer-F & gRNA a Primer R

2 – gRNA a Primer F & gRNA b Primer R

3 - gRNA b Primer F & gRNA c Primer R

Use the pGTR plasmid as template

-- --- --- ---- ---- ---- ----

An example:

G-primer-F

5’ AGgaagacTACTCGAACAAAGCACCAGTGG 3’

gRNA14

Forward sequence: 5’ TTGCAGCAGTGGAGAAAAGA 3’

Reverse complement sequence: 5’ TCTTTTCTCCACTGCTGCAA 3’

GLK14-a-F

5’ GAgaagacATGTGGAGAAAAGAGTTTTAGAGCTAGAA 3’

GLK14-a-R

5’ GAgaagacTACCACTGCTGCAATGCACCAGCCGGGAA 3’

gRNA3

Forward sequence: 5’ GGGCGAGGGCTTCAAGATAC 3’

Reverse complement sequence: 5’ GTATCTTGAAGCCCTCGCCC 3’

GLK3-b-F

5’ GAgaagacATGCTTCAAGATACGTTTTAGAGCTAGAA 3’

GLK3-b-R

5’ GAgaagacTAAAGCCCTCGCCCTGCACCAGCCGGGAA 3’

gRNA7

Forward sequence: 5’ AGGATATGGAGTGGGTTGCT 3’

Reverse complement sequence: 5’ AGCAACCCACTCCATATCCT 3’

Final-c-GLK7

5’ GAgaagacTATAAAACAGCAACCCACTCCATATCCTTGCACCAGCCGGGAATC 3’

Amplify using PCR from pGTR plasmid, Gel extract and then directly clone into L1 plasmids using BbsI GG reaction

For the PCR, use Phusion (Thermo) or a similar proofreading polymerase

PCR cycling conditions were: 98^o^C (1:30), [98^o^C (30sec), 55^o^C (30sec), 72^o^C (45sec)] x35, 72^o^C (5min)

Primer combination is:

1 - G-primer-F & GLK3-b-R

2 – GLK3-a-F & GLK14-b-R

3 - GLK14-b-F & Final-c-GLK17

**B- Loop assembly cloning of tRNA-gRNA parts into the L1 vector to combine tRNA-gRNA parts with the Mp*U6* promoter**

Plasmid concentrations should be according to (Sauret-Güeto et al. 2020). Aliquots of the DNA part were prepared at a concentration of 15 nM and of the acceptor vector at a concentration of 7.5 nM

To calculate the concentration in ng/μL:

- For a final concentration of 15 nM, the concentration in [ng/μL] equals N (the length in bp of the plasmid) divided by 110. This is an approximation of the formula:

15∙10^(-9)mol/L x ((607.4 x N ) + 157.9)g/mol x 10^(-6)L/μL x 10^9ng/g = concentration (ng/μL)

- For a final concentration of 7.5 nM, the concentration in [ng/μL] equals N divided by 220.

Prepare reaction master mix (in μL):

| MilliQ H_2_O | Up to 20 |
| --- | --- |
| BSA (1 mg/mL) | 1.5 |
| 10x T4 DNA Ligase buffer (NEB) | 2 |
| T4 DNA Ligase at 400 U/μL (NEB, #M0202S) | 1.5 |
| BbsI-HF 10 U/μL (NEB, #R3539) | 1.5 |
| OP-074 or OP-075 plasmids | 1 |
| tRNA-gRNA parts (Gel extracted) | 1 μL per part |

-Place samples on the thermocycler and incubate using the following program: Loop Assembly: [3 minutes at 37^o^C and 4 minutes at 16^o^C] x26, Termination: 5 minutes at 50^o^C and 10 minutes at 80^o^C

-Transform chemically competent using 7-10 μL of reaction and plate on LB agar plates with 50 μg/mL kanamycin and 40 μg/mL X-gal. Incubate at 37^o^C for 16 h.

-Confirm with Sanger sequencing

**C- Loop assembly cloning into the L2 pCsA acceptor to combine of tRNA-gRNA with the transcription unit for Cas9 expression**

Prepare reaction master mix (in μL), plasmid concentrations should be as above and according to (Sauret-Güeto et al. 2020):

| MilliQ H_2_O | Up to 20 |
| --- | --- |
| 10x T4 DNA Ligase buffer (NEB) | 2 |
| T4 DNA Ligase at 400 U/μL (NEB, #M0202S) | 1.5 |
| SapI 10 U/μL (NEB, #R0569S) | 1.5 |
| OP-074 or/and OP-075 plasmids (or appropriate pCk spacers) | 1 each |
| Selection marker plasmid | 1 |
| pCsA | 1 |

-Place samples on the thermocycler and incubate using the following program: Loop Assembly: [3 minutes at 37^o^C and 4 minutes at 16^o^C] x26, Termination: 5 minutes at 50^o^C and 10 minutes at 80^o^C

-Transform chemically competent using 10-12 μL of reaction and plate on LB agar plates with 50 μg/mL spectinomycin and 40 μg/mL X-gal. Incubate at 37^o^C for 16 h.

-Confirm with Sanger sequencing

After cloning in the L2 acceptor the tRNA-gRNA construct will be:

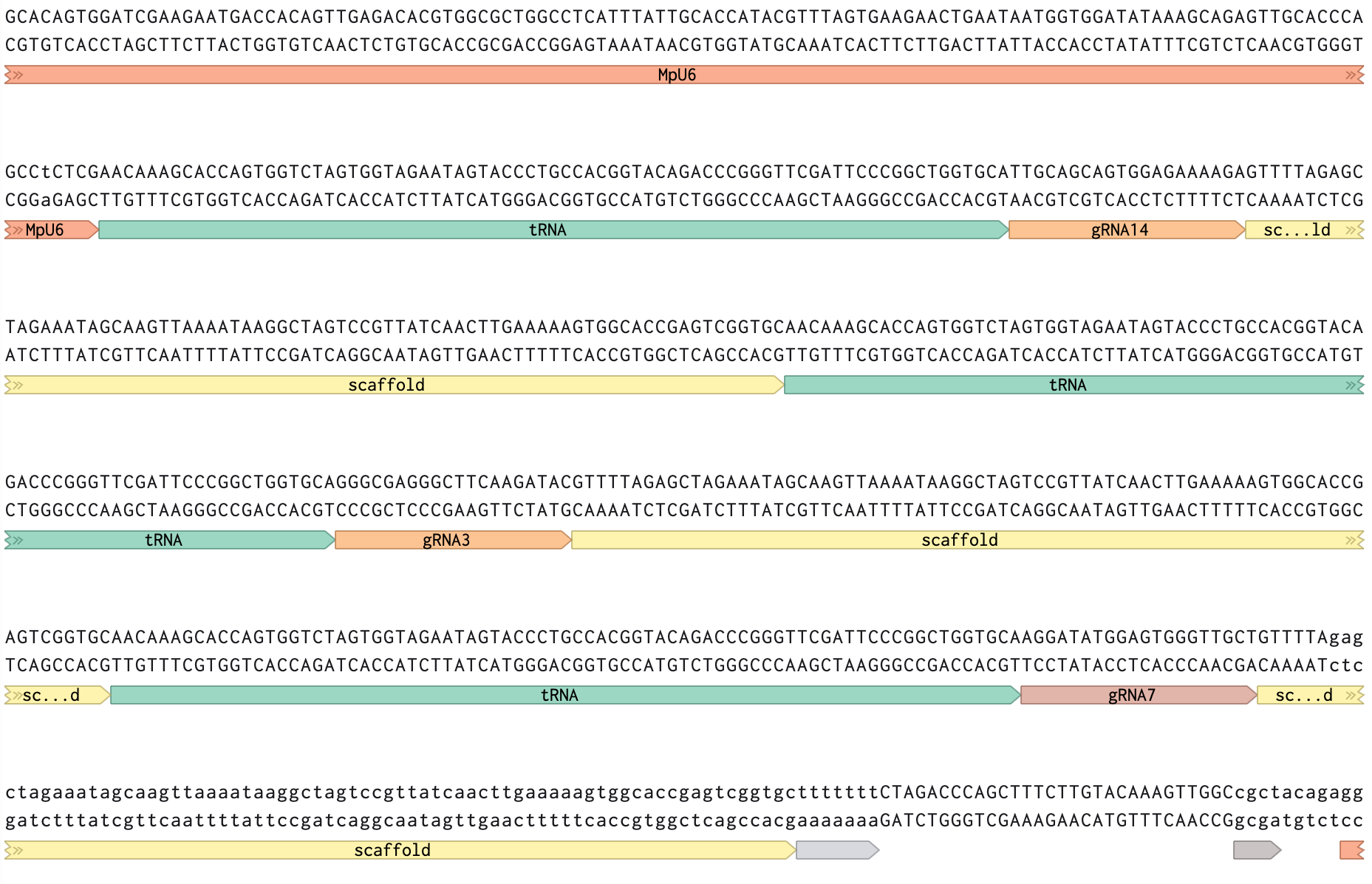

For the plasmid map of the final construct see file: “l2-glk-3g-csa.gb”
